# Extending Specimens to save Plant DNA: structuring Department DNA collections in times of Biodiversity loss

**DOI:** 10.1101/2025.05.20.655165

**Authors:** Claudia González-Toral, Eduardo Cires

**Author notes:** Corresponding author: Claudia González-Toral: e-mail address /; Eduardo Cires.

## Abstract

Plant biodiversity DNA banks are scarce despite the current plant biodiversity loss, their value for *ex situ* conservation, DNA preservation progress and the rapid growth of DNA-dependent research fields. We explore the principles and basic organization of plant biodiversity DNA banking and propose new ways of addressing the biodiversity loss through their implementation. Small Department collections could be created through a 6-step holistic process aiming to interconnect 3 types of collections (DNA extracts, DNA-rich tissues and herbarium vouchers) while generating Extended Specimens that would contribute to local and global plant biodiversity knowledge and conservation efforts. We propose a change in the international biobanking strategy (the Dynamo scheme) as interconnecting many small Department collections would put fewer samples at risk in case of catastrophe, optimise sampling strategies and cover in depth more taxa and distribution ranges, while encouraging national and international collaborations.

## Introduction

Biological collections surged three centuries ago, becoming fundamental for scientific advances in fields like medicine, taxonomy, ecology, evolutionary biology or genetics (Pyke & Ehrlich, 2010; Holetschek *et al.,* 2012; Kang *et al.,* 2013; Martin & Kaye, 2000). Biobanks—research-oriented facilities for long-term storage of collections of well-preserved samples and their associated data—appeared in 1948 in the United States of America (USA) when a human specimens collection was systematised into the Framingham Heart Study (FHS) blood bank (Thormann *et al.,* 2006; Kang *et al.,* 2013; Parodi, 2015). The advances in preservation technologies led to the transformation of rudimentary human specimens collections into biobanks (Kang *et al.,* 2013; Souza & Greenspan, 2013), resulting in the establishment of many biobanks types during the following decades (e. g. tissue, disease-specific or population specific biobanks) (Parodi, 2015).

During the late 1960s, international organizations like the Food and Agriculture Organization of the United Nations (FAO) regarded the loss of animal and plant genetic diversity as a potential conservational problem, drawing the attention of scientific institutions towards genetic biorepositories during the following two decades, due to their potential as ex situ conservation tool for genetic diversity (Wildt *et al.,* 1993; Heywood, 1995; Thormann *et al.,* 2006; León-Lobos *et al*., 2012; Seberg *et al.,* 2016; Singh & Singh, 2017). This prompted the appearance of animal and plant germplasm collections associated to biodiversity-centred institutions (e. g. museums, zoos, botanic gardens and university departments) aiming to mitigate the biodiversity loss due to anthropic activities (also known as the Biological Diversity Crisis (BDC)) and ensure populations and species future viability (Wilson, 1985; Wildt *et al.,* 1993; Krauss *et al.,* 2002; Palacio-Mejía, 2006; Souza & Greenspan, 2013; Leigh *et al.,* 2019). Collections evolved into systematised biobanks preserving various types of high-quality organic material and their associated information: germplasm banks (explants storage), seed banks, gene banks (vegetative propagules), tissue banks, pollen banks or DNA banks (Wildt *et al.,* 1993; Adams, 1997; Palacio-Mejía, 2006; Rice, Henry & Rossetto, 2006; Thormann *et al.,* 2006; Astrin *et al.,* 2013; Rajasekharan *et al.,* 2013; Souza & Greenspan, 2013; Dröge *et al.,* 2016; Comizzoli & Wildt, 2017; Ruta *et al.,* 2020).

Biodiversity DNA biobanks focus on wild and/or commercial species complete other collections (e. g. universities, Botanic Gardens) and may hold DNA libraries, cloned fragments or complete plant genomes (Organisation for Economic Co-operation and Development (OECD), 2001; Rice, Shepherd *et al.,* 2006; Jain *et al.,* 2012; Droege *et al.,* 2014; Watson, 2014; International and Society for Biological and Environmental Repositories (ISBER), 2018).

The 1990s and 2000s DNA technology advances and costs reductions allowed extending the use of DNA sequences for phylogenetic and genetic diversity studies, prompting the appearance of department private DNA and/or DNA-rich plant tissue collections associated to ecological data (Adams, 1997; Karp *et al.,* 1997; Jenkins, 2003; Rice, Shepherd *et al.,* 2006; Palacio-Mejía, 2006; Gaudeul & Rouhan, 2013; Shaw *et al.,* 2014; Watson, 2014; Rabeler, 2017). These department private DNA collections were the precursors of the modern DNA Plant Biodiversity Banks or Biobanks (PBBs), institutions specialised in the long-term storage and preservation of high-quality DNA samples (DNA extracts, DNA-rich tissue material or both) and theirs associated information, which are fundamental tools for addressing the botanical problems arisen from the BDC (Blackmore, 2002; Thormann *et al.,* 2006; Hodkinson *et al.,* 2007; Astrin *et al.,* 2013; Gaudeul & Rouhan, 2013; Dröge *et al.,* 2016).

Nevertheless, despite of prompting accessibility to high-quality and properly managed genomic samples, complementing conservation strategies and the 2000s claims and estimations of the proportionally low costs of establishing DNA biobanks at global scale, PBBs have not been fully integrated in the already existing biodiversity preservation facilities (Savolainen, 2004; Hodkinson *et al.,* 2007; Brown, 2011; Astrin *et al.,* 2013).

The BDC has aggravated during the last four decades due to The intensification of processes directly associated to high extinction rates of animals, plants and fungi, habitat loss and fragmentation has aggravated the BDC during the last decades(e.g. Peres, 2001; Mortelliti *et al.,* 2011), introduction of alien species with invasive potential (e.g. Manenti *et al.,* 2018; Muthukrishnan & Larkin, 2020), species overexploitation (e.g. Bodeker *et al.,* 2014; Chuanwu *et al.,* 2019), changes in land use and Climate Change (e.g. Sekercioglu *et al.,* 2008; Feeley & Silman, 2010; De Baan *et al.,* 2013) (De Vos *et al.,* 2015; Eichenberg *et al.,* 2021; International Union for Conservation of Nature (IUCN), 2024). These synergetic processes are boosting the extinction rates by 1,000 times the background rates, with expectations of reaching 10,000 times in the future and no signs of deceleration due to financial crises (De Vos *et al.,* 2015; Steffen *et al.,* 2015). The Intergovernmental Panel on Climate Change (IPCC) projections for terrestrial and freshwater ecosystems are daunting, as estimations suggest a global mean temperature raise of 2-4°C, a 35-40% and 15-35% increase of wildfires and biome shifts while a 5°C increase would trigger the extinction of 60% of the species inhabiting the affected ecosystems (Parmesan *et al.,* 2022). Currently, 129 plant species have gone completely extinct, 45 are extinct in the wild and over 320,000 flowering plant species are comprised in IUCN Red List of Threatened Species, in a context were 77% of the estimated 100,000 plant species unknown to science are estimated to be under extinction threat and global risk assessments have not reached 45% of the threatened red-listed vascular plants (Nic Lughadha *et al.,* 2020; Antonelli *et al.,* 2023; IUCN, 2024). Furthermore, other factors like pollinators abundance decrease could accelerate plants decline, something with potentially devastating and with unforeseeable consequences for ecosystems given plants’ fundamental ecosystemic role (Grime, 2002; Schleuning *et al.,* 2016; Traveset *et al.,* 2017). BDC and Climate Change interact synergistically and have economic, social and ecological consequences at global scale; thus becoming main concerns for biodiversity-focused research (Mace *et al.,* 2014; Urban, 2015; Román-Palacios & Wiens, 2020; Kedward *et al.,* 2023; Schlaepfer *et al.,* 2023).

The biodiversity loss statistics and risk assessments do not usually take into account the genetic dimension, a relevant aspect for species and population survival as genetic diversity loss is intimately related to extinction debt, which is one of the main issues for plant extinctions estimations (Helm *et al.,* 2009; Garner *et al.,* 2020; Nic Lughadha *et al.,* 2020). Unfortunately, the deficit in plant biosiversity biobanking (PB biobanking) reported by Hodkinson *et al*. (2007) is still happening in a context were biodiversity loss poses an ecological and financial threat to human society, plant extinctions have unpredictable ecosystemic effects, global risk assessment are insufficient and tend not to include genetic data, and there is a chronic insufficient funding for non-commercial plants research and conservation (Suarez & Tsutsui, 2004; Bradley *et al.,* 2014; Urban, 2015; Schleuning *et al.,* 2016; Roberson & Meyer, 2020; Kedward *et al.,* 2023).

Moreover, despite its potential for boosting basic science and plant conservation, PB biobanking principles have not been fully implemented within the existing biological collections (Hodkinson *et al.,* 2007; Astrin *et al.,* 2013; Gaudeul & Rouhan, 2013; Droege *et al.,* 2014; González-Toral & Cires, 2022). In this context, we aim to explore (1) how PB biobanking principles and activities meet the current scientific and social goals for biodiversity presrvation and information sharing and (2) new ways in which the scientific community can address the biodiversity loss through the implementation of PB biobanking principles.

## Methods

We conducted a search of English and Spanish language scientific papers and books in “Google Scholar” (https://scholar.google.com/) and “Web of Science” (https://webofscience.com/wos/) from March 2020 to May 2024. We used the key words: “Biobank”, “DNA Bank”, “Plant Biodiversity”, “DNA preservation”, “biobanking”, “Nagoya Protocol”, “Access and Benefit-Sharing”, “Biological Diversity Crisis” and “Plant Conservation”.

### PBBs structure, functioning and classic global scheme

PBBs are instrumental for the Nagoya Protocol on Access to Genetic Resources and the Fair and Equitable Sharing of Benefits Arising from their Utilization (= Nagoya Protocol) implementation, complementing conservation strategies, providing accessibility to properly-managed genomic samples, and generating new phylogenetic knowledge (Blackmore, 2002; Hodkinson *et al.,* 2007; Secretariat of the Convention on Biological Diversity (CBD), 2010; Astrin *et al.,* 2013; Droege *et al.,* 2014; Consortium of Europena Taxonomic Facilities (CETAF), 2020). This is accomplished thanks to their structure and functioning, which are based on a series of “Major Operational Activities” (MOAs) implemented through “standardized operating procedures” (SOPs) (Adams & Adams, 1992; Hodkinson *et al.,* 2007; Astrin *et al.,* 2013; Watson, 2014). PB biobanking usually involve 5 MOAs—(1) material collection, (2) storage of the plant tissue, (3) DNA extraction, (4) DNA banking and (5) DNA utilisation—integrated in a two-node structure consisting in a Working Node (WN) and a Reserve Node (RN) (Adams & Adams, 1992; Adams, 1997; Rice, Shepherd *et al.,* 2006; Hodkinson *et al.,* 2007; Astrin *et al.,* 2013; Watson, 2014; Zimkus & Ford, 2014a) (Figure 1). The WN is responsible for sample collection and acquisition, DNA extraction, banking and utilization under controlled conditions (i. e. loans, shipments and interactions with researchers and institutions); meaning that its SOPs are directly related to the Nagoya Protocol implementation (Adams & Adams, 1992; Mattick, Ablett & Edmonson, 1992; Adams, 1997; Rice, Henry *et al.,* 2006; Hodkinson *et al.,* 2007). The RN focuses mainly on tissue banking, usually acts as a backup for the WN and/or for other biobanks with different scopes (Adams & Adams, 1992; Adams, 1997; Caujapé-Castells *et al.,* 2006; Hodkinson *et al.,* 2007; Zimkus & Ford, 2014a; South African National Biodiversity Institute (SANBI), 2020). Samples and metadata are connected through an effective labelling system to secure specimens’ traceability (Global Genome Biodiversity Network (GGBN), 2015; Funk *et al.,* 2017; ISBER, 2018; CETAF, 2020).

**Figure 1.**
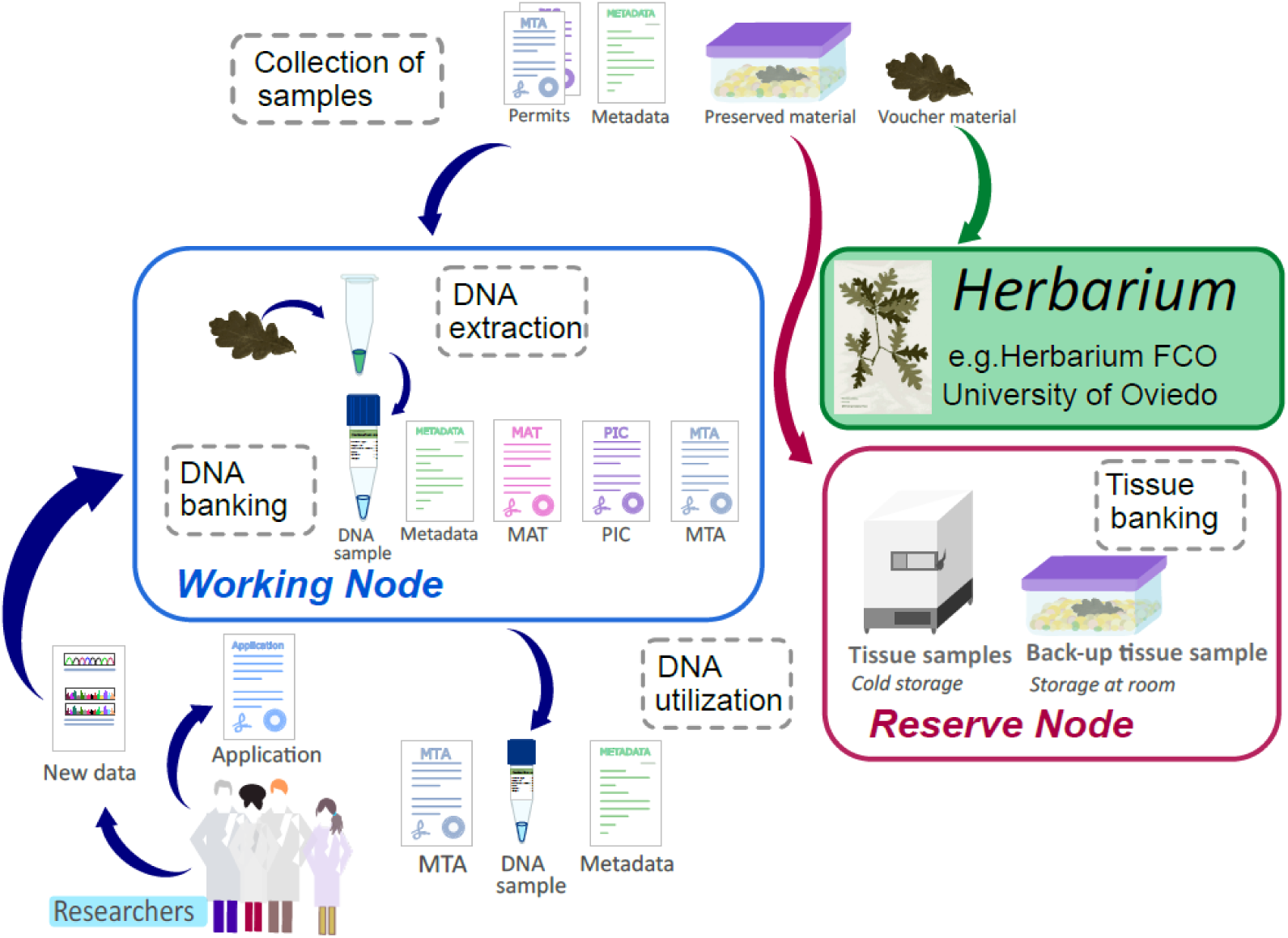
Scheme depicting the main Major Operational Activities (MOAs) of a plant DNA biobanks described as described by Hodkinson *et al.,* (2007) and the main functions of the working and reserved nodes as proposed by Adams (1997) for the DNA bank-Net. The main MOAs correspond to the name in the black cage, while the working node and the reserve node correspond to the blue and red cages respectably.

Preservation techniques improvements resulted in the establishment of various types of plant biobanks storing different tissues: explants under controlled sterile conditions for slow or suspended growth (germplasm biobanks), viable seeds of crops and their wild relative (e.g. Millennium seed bank), vegetative propagules (gene banks), different types of tissue (e. g. Alexander von Humboldt Institute (AvHI) tissue bank), pollen (e.g. the pollen collection of the ICAR-National Bureau of Plant Genetic Resources (ICAR- NBPGR) of India) or DNA extracts (e.g. Kew Royal Botanic Garden) (Palacio-Mejía, 2006; Thormann *et al.,* 2006; Rajasekharan *et al.,* 2013; Ruta *et al.,* 2020). PBBs may focus on certain type of organisms, on a specific scientific sampling strategy or even have specific geographic scope. For instance, the Japanese National Institute of Agrobiological Sciences (NIAS) Genebank (https://www.gene.affrc.go.jp/about_en.php) focuses on agricultural taxa and their wild relatives, while Kew DNA Bank (https://dnabank.science.kew.org/homepage.html) gather samples from wild species (NIAS, 2024). The Kostrzyca Forest Gene Bank (https://www.lasy.gov.pl/en/information/news/kostrzyca-forest-gene-bank-collects-dna-of-endangered-plants-in-the-bialowieza-forest) gathers samples from the eastern European Białowieża Forest, The Pacific Center for Molecular Biodiversity (PCMB) (https://www.bishopmuseum.org/pcmb/) focuses on a concrete bioclimatic region, while Kew DNA Bank (https://dnabank.science.kew.org/homepage.html) has a wider scope and accepts samples from all over the world (Campbell *et al.,*, 2018a). In other cases, eDNA collections like the Chicago Botanical Garden’s DNA Biorepository (https://www.chicagobotanic.org/) comprise several individuals from each taxa, making them adequate for phylogenetic and taxonomic purposes.

In the recent years, the relevance of physical vouchers for DNA-based studies has been highlighted (e. g. Dick & Webb, 2012; Culley, 2013; Gostel *et al.,* 2016; Buckner *et al.,* 2021), leading to the appearance of new terms referring to their relationship to genetic samples (Pleijel *et al.,* 2008; Funk *et al.,* 2018) (Table 1) or to their interconnection to all their natural and derived information (Webster, 2017; Lendemer *et al.,* 2020). These theoretical frameworks sustain that vouchers and genetic samples should ideally be the same individual—i. e. hologenophore direct voucher or, slightly suboptimally, a isogenophore direct voucher (Table 1)—, that besides phenotype and genotype, the ecological context should be preserved through metadata creating Extended Specimens (ESs); and that all direct and derived samples (e. g. genetic samples or histologic samples) and their data (e. g. site of collection and DNA sequence or histologic images) should be connected for a better understanding of species and populations, generating Extended Specimens Networks (ESNs) (Pleijel *et al.,* 2008; Webster, 2017; Funk *et al.,* 2018; Lendemer *et al.,* 2020). PBBs internal functioning generate *de facto* a collection of ESs and ESNs (mainly hologenophore direct vouchers) preserved under controlled conditions and available for researchers, something that contributes to the reproducibility of results without taxa resampling (GGBN, 2015; Funk *et al.,* 2017; ISBER, 2018; CETAF, 2020). Besides, PBBs can contribute to natural collections by promoting large expeditions in collaboration with specialists from museums and botanic gardens—reducing the costs of plant biodiversity DNA-based studies— and to scientific studies by providing services to researchers unfamiliar with molecular techniques and by generating MOAs and SOPs protocols’ rankings (Mattick *et al.,* 1992; Palacio- Mejía, 2006; Hodkinson *et al.,* 2007; Zimkus & Ford, 2014a, 2014b; Spooner & Ruess, 2014a, 2014b; Lear *et al.,* 2018; ISBER, 2018; Abdaljaleel *et al.,* 2019; SANBI, 2020; CETAF, 2020). The reliability and low costs of DNA-rich tissues preservation in silica gel allows sparing morphological voucher destruction (Staats *et al.,* 2011; Gaudeul & Rouhan, 2013; González-Toral *et al.,* 2021) and represents a great opportunity to compare its basal decay with that of DNA extracts preserved under controlled conditions.

**Table 1.**
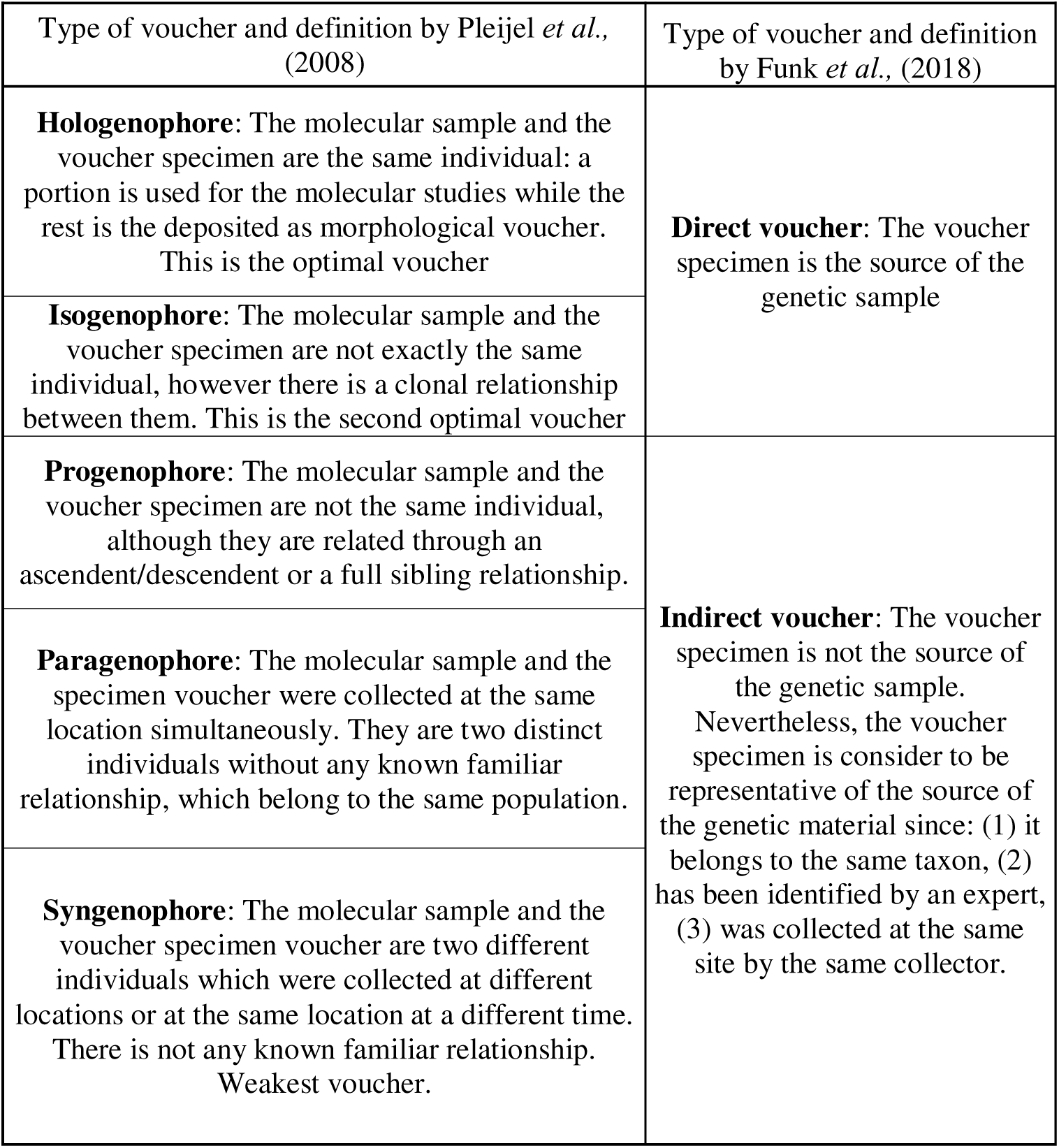
Types of morphological voucher regarding its relation to the samples of molecular studies and their taxonomic value described by Pleijel et al. (2008) and by Funk et al. (2018).

The BDC triggered the appearance of regulatory international legislation for biodiversity, wild life and genetic resources such as the Convention on International Trade in Endangered Species of Wild Fauna and Flora (CITES) (IUCN, 1973), the International Treaty on Plant Genetic Resources for Food and Agriculture (ITPGRFA) (FAO, 2001) and the Convention on Biological Diversity (CBD) and the specifications of the Nagoya Protocol (CBD, 2010). Since expeditions, donations and samples exchanges are essential parts of biobanks’ material collection and DNA utilisation MOAs, PBBs activities makes them play an important role in the implementation of these international legislations (Droege *et al.,* 2014; Zimkus & Ford, 2014a, 2014b; Campbell *et al.,* 2018; CETAF, 2020). The CBD and the Nagoya Protocol (ratified by 126 countries) are the most relevant international legislations for PBBs, as they guarantee equality and justice by clarifying genetic resources’ accessing and benefit-sharing terms (CBD, 2010; GGBN, 2015; Biber-Klemm *et al.,* 2016; Davis & Borisenko, 2017). Nagoya Protocol elaborated the CBD principles for the Access and Benefit-Sharing (ABS) through Prior Informed Consents (PICs), which abides the local legislation of the community holding the sovereign right over the sample and specifies the accessibility terms, and Mutually Agreed Terms (MATs), which unambiguously defines provider’s monitoring process and usually includes genetic material and traditional knowledge accessibility, publishing and storage policies (GGBN, 2015; Biber-Klemm *et al.,* 2016; Davis & Borisenko, 2017; ISBER, 2018). Once the PICs and MATs for the ABS are reached, its information is published in the Access and Benefit- sharing Clearing-House (https://absch.cbd.int/) by the provider’s national competent authority to generate an Internationally Recognized Certificate of Compliance (IRCC) number, which should always be associated to samples (Davis & Borisenko, 2017). The PICs and MATs terms have to be reflected in the biobank’s Material Transfer Agreements (MTAs) when loaning samples in compliance with the CBD and the Nagoya Protocol (Rice, Shepherd *et al.,* 2006; Zimkus & Ford, 2014a; Biber-Klemm *et al.,* 2016; Davis & Borisenko, 2017) (Figure 2). All ABS agreements require: (1) internal protocols for establishing relationships with the provider’s competent authorities and implementing their policies, (2) settling technical approaches and the potential extent of scientific research and (3) determining the benefits-sharing terms, usually not commercial for PBBs (e. g. scientific training or future collaborations) (GGBN, 2015; Davis & Borisenko, 2017; Campbell *et al.,* 2018; ISBER, 2018; Williams, 2019; CETAF, 2020).

**Figure 2.**
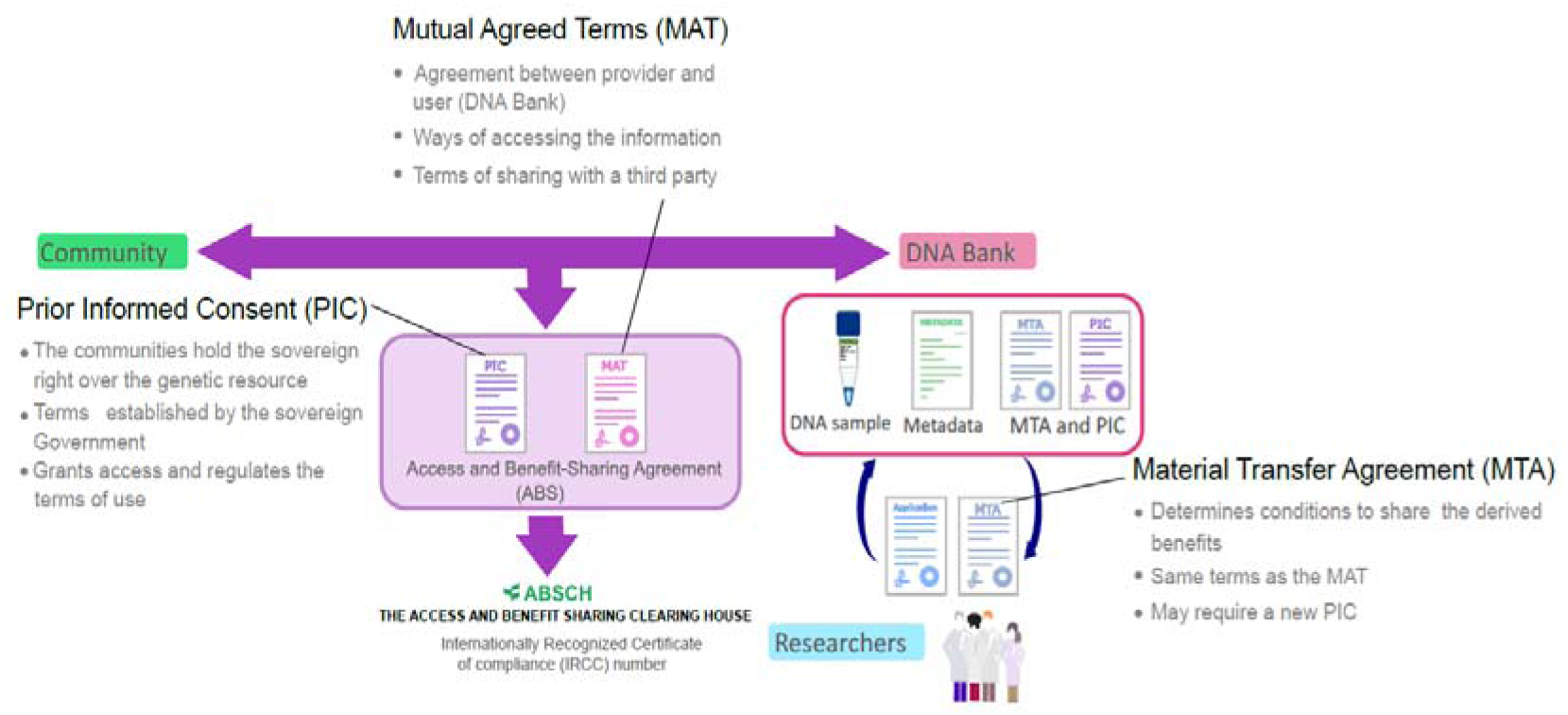
Scheme representing the different agreements required by the Nagoya Protocol for acquiring, donating and loaning samples to a DNA bank while conveying with the international legislation.

Therefore, PBBs are fundamental for generating and sharing tissue and DNA preservation SOPs, the creation of ESs and ESNs and the implementation of national and international legislation for plant biodiversity preservation.

### Changing the scheme of PBBs networks

The idea of creating a network that would facilitate the connection between the existing biobanks emerged in the early days of DNA biobanking, being DNA-Bank Net— an ambitious project aiming to preserve endangered tropical plants DNA in all continents— the first initiative of this kind (Adams 1988, 1993; Spooner & Ruess, 2014a). Initially, few scientists expressed interest however, with time, its scientific interest grew so that by the early 1990s scientists from 40 institutions from all continents were compelled, leading to a 1991 meeting in which the nodes-structure and the idea of physically separating RN (or ideally the two RNs) from the WN by harbouring them in the facilities of DNA banks found at different continents were proposed (Adams & Adams, 1992; Adams 1993, 1997; Graner *et al.,* 2006; Spooner & Ruess, 2014a). Despite its useful contributions and the 2000s estimations of the proportionally low costs of establishing DNA biobanks at global scale, DNA-Bank Net remained inactive by the mid-2000s (Savolainen, 2004; Graner *et al.,* 2006; Hodkinson *et al.,* 2007; Brown, 2011; Spooner & Ruess, 2014a). Furthermore, during these decades, public funding of taxonomic and systematics research declined up to a point in which the existing specimens’ collections generated during decades were threatened, despite of their many social and scientific benefits (Dalton, 2003; Suarez & Tsutsui, 2004).

Later on, the creation of a network of biodiversity biobanks was reassumed by the decentralised pilot project DNA Bank Network, in which all types of biodiversity independent DNA collections coordinated to make their data accessible through the internet and generate a larger and more diverse pool of vouchers-linked sequences (Gemeinholzer *et al.,* 2011). In 2011, the DNA Bank Network evolved into the Global Genome Biodiversity Network (GGBN) an association that aims to standardize sharing procedures, ethical use and best practices while encouraging knowledge exchange and recruiting new members (Droege *et al.,* 2014; Seberg *et al*., 2016; GGBN, 2024). The GGBN has been growing in the recent years: from 24 members in 2014 to 112 biorepositories from 38 countries, including at least 29 plant and fungi collections and 2 department collections (Dröge, *et al.,* 2014; GGBN, 2024). A part from providing guides and SOPs implementation models and the ABS, GGBN’s compromise with the Tree of Life led to the creation of the GGBN data portal, which addressed the problem of DNA banks samples information availability by allowing taxa and samples searches within the members’ standing collections (having over 185,000 accessions from Kingdom Plantae)— and by creating data standards (e. g. Sample PREanalytical Code (SPREC) or ABCD-DNA) (Benson *et al.,* 2010, 2011; Betsou *et al.,* 2010; Dröge *et al.,* 2014, 2016; Seberg *et al*., 2016; GGBN, 2024). However, despite the GGBN globalization efforts, most members are not located within the highest plant biodiversity richness regions (Barthlott *et al.,* 2005), but rather in western richer countries (GGBN, 2024). On the other hand, there is no information on whether GGBN follows the RN relocation scheme of Adams (1997).

The Adams (1997) RN relocation scheme was based on the idea of preserving plant biodiversity by securing biobanks samples from all sorts of threats, including natural disasters. As one of the main aims of DNA-Bank Net was to preserve as many species as possible at reduced costs, it focused on large collections (i. e. herbaria and botanic gardens) since their personnel is already familiarised with sampling SOPs (Adams & Adams, 1992; Adams, 1997). However, although this was a well-thought scheme that addressed the main problems of plant biodiversity loss using large collections as backbone, it has not been implemented. Moreover, during the 1990s and 2000s, when the scientific community and the society were aware of the BDC, biodiversity collections faced founding reductions or even closures that have tremendously affected the scientific value of the remaining collections (Adams & Adams, 1992; Dalton, 2003; Lavoie, 2013). Hence, the success of using large collections as backbone for preserving plant biodiversity depends, in part, on funding, the potential financial crises and political will. Since the Adams (1997) scheme has not been implemented worldwide (although there exist examples like the South African National Biodiversity Institute (SANBI) (SANBI, 2020)), any disaster at any large collection would represent the loss of a large proportion of genetic information in a context were world biodiversity is on decline (Eichenberg *et al.,* 2021).

It should also be highlighted that smaller DNA collections (i. e. Department collections) would be of interest for meeting the goal of preserving genetic material and metadata of as many species as possible. These collections have many appealing features to fulfil this task as they may (1) focus on a specific biogeographic unit and their related areas and/or on specific groups of families, (2) present samples from species, subspecies or populations of conservational value (e. g. endemisms, relictic populations), (3) have associated morphological voucher material deposited in Index Herbariorum listed Herbaria, (4) many of the data derived from those samples has been published and is publically available (e. g. Genbank sequences. https://www.ncbi.nlm.nih.gov/genbank/) and (5) samples are usually preserved as tissues in silica gel and/or as DNA extracts in cold. For example, our DNA collection at the area of Botany of the department of Biology of Organisms and Systems of the University of Oviedo is formed by DNA extracts preserved at -20°C and foliar tissue preserved in silica gel at room temperature from taxa and populations of taxonomic and conservational interest (e. g. Cires Rodríguez, Cuesta Moliner & Fernández Prieto, 2012; Cires & Prieto, 2015; González-Toral *et al.,* 2023) occurring in the Mountains Cantabrian and in the European Atlantic Biogeographic unit and others related to it (e. g. Mediterranean) and associated to morphological herbarium vouchers, most of them deposited in the Index Herbariorum listed Herbario de la Facultad de Ciencias de la Universidad de Oviedo (FCO) (Index Herbariorum, 2023; Universidad de Oviedo, 2023). These two ways of preserving DNA and DNA-rich tissues represent an opportunity for following the Adams & Adams (1992) and Adams (1997) two-node scheme from an early stage, as the WN could be formed by the DNA extracts and the RN could be built from the tissues preserved at room temperature. Furthermore, the MOAs could have a holistic approach focused not only on generating ESs (Webster, 2017), but also on ESNs (Lendemer *et al.,* 2020) by connecting the published data with the curated specimens (i. e. DNA and DNA-rich samples with DNA sequences and herbarium vouchers).

Given their potential, smaller Department DNA collections like ours may have an important role to play in plant biodiversity conservation as the combination of many of these collections could (1) cover greater areas of floras and biogeographic regions than the larger existing collections, (2) cover lower taxonomic levels (e. g. subspecies, *formas*) and (3) focus on the genetic diversity of populations with local and/or regional conservational interest. Besides, given the critical funding-dependence of large collections witnessed in the last decades and the chronical underfunding of plant conversation programs (Suarez & Tsutsui, 2004; Balding & Williams, 2016; Roberson & Meyer, 2020), the existence and maintenance of many smaller collections would secure samples survival. We believe that the current situation of PB biobanking is analogous to the 1940 Dunkirk evacuation (also known as Operation Dynamo): having a small number of larger collections as suggested by Adams (1997) allows for the preservation of many species at one place; however, this put at risk many samples at once if there was any catastrophe (including financial) to happen. On the other hand, having a large number of smaller collections requires many more institutions involved and a higher level of coordination but, allows to preserve substratially more samples and puts at risk a smaller number of samples in case of catastrophe. Thus, this “Dynamo scheme” is based on the existence of many interconnected smaller collections that could all in all preserve more samples from taxa and populations. Therefore, although the current number of Department DNA collections is unknown, they may be of vital importance in future scenarios of Climate Change, species extinctions and population and species loss of genetic diversity.

### Developing a department PBB from tissue and DNA collections

Given all the features and potential of department collections, we present a 6-step process for creating a small PBB focused on a specific floristic area and on generating ESs and ESNs (Figure 3). These phases aim to set the PBB’s structure and functioning and to gather and assess the current knowledge of its target floristic area, while gradually cataloguing specimens and making them available. As we pursue a holistic approach that facilitates information sharing and the Dynamo scheme, collaborating with existing PBBs and digitalizing and centralising specimens’ open-access information will be a fundamental to this process.

**Figure 3.**
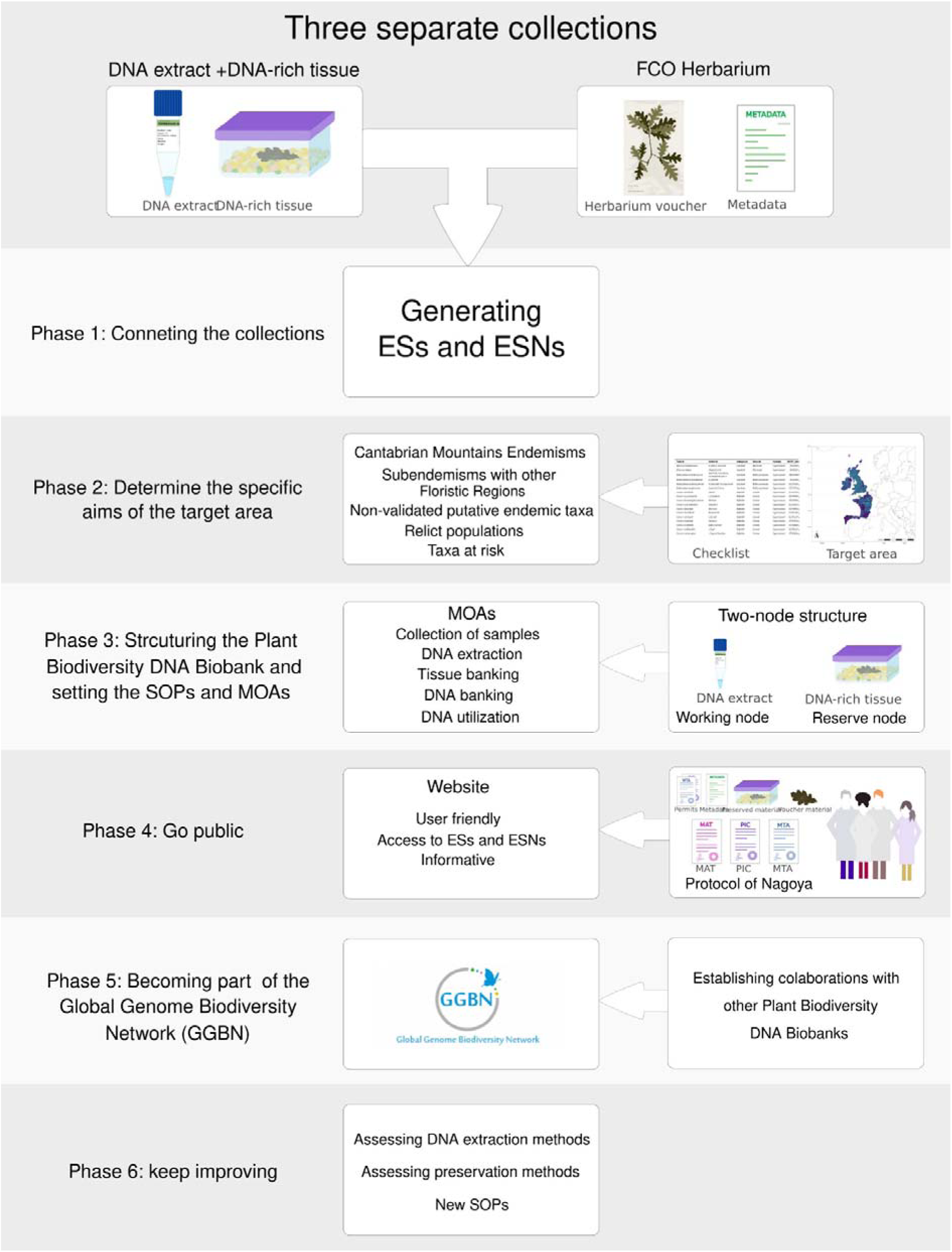
Scheme depicting the different phases of the process for transforming 3 separate collections into a Department Plant Biodiveristy DNA biobank using the collections of the area of Botany of the department of Biology of Organisms and Systems of the University of Oviedo as example.

### Phase 1: Connecting DNA collections to herbarium vouchers and metadata: creating an ESs collection

Initially, the DNA collection specimens (e. g. the DNA extract and/or the DNA-tissue samples) should be connected to their herbarium vouchers and metadata, their publically-available genetic information (e. g. Genbank ID number) and the published scientific works from in which they derive. This is essential for creating ESs and ESNs and implies generating an inventory of published information associated to each sample.

Thus, a private database connecting the three separate collections samples (or two collections if there is no tissue collection) and physically connecting the specimens via ID numbers and barcodes or QR codes would be the PBB basis. Metadata deficiencies detected during this connection process would help to polish the data gathering process, impacting positively on SOPs, ESs, ESNs and FAIR data and implementing the Responsible Research and Innovation (RRI) principles (Hodkinson *et al.,* 2007; Owen, Macnaghten, & Stilgoe, 2012; Funk *et al.,* 2017; Williams *et al.,* 2020).

Three fundamental aspects should be taken into account during this phase: (1) taxa names update and standardization, (2) metadata gathering and standardization and (3) sampling strategy. The first aspect could be solved by following the criteria of World Checklist of Vascular Plants (WCVP) and World Flora Online (WFO) (Govaerts *et al.,*, 2021; Royal Botanic Gardens Kew, 2023;WFO, 2023) during an initial state and then modifying it according to the latest literature. The DNA collection’s specimens initial database should ideally include: (1) taxon name, (2) taxonomy (i.e. kingdom, division, class, order), (3) accepted genus (including author(s)), (3) species accepted (including author(s)), (4) infraspecific accepted taxon, if applicable, (5) accepted species WCPV number and its url, (6) location (including coordinates and altitude), (7) collection date, (8) collector(s), (9) voucher ID number, (10) DNA extract and/or tissue sample ID, (11) type of study conducted on samples (i. e. phylogenetic vs genetic diversity), (12) molecular information obtained from the sample (i. e. sequenced markers or genomic sequencing), (13) sample’s National Center for Biotechnology Information (NCBI) ID (i.e. GenBank number), if applicable and (14) publication (e. g. Title, doi) (Table 2). The future acquisitions would benefit from the addition ecological data during the collection process, including generating photovouchers as nowadays they can be easily obtained with cell phones. Special attention should be given to the recommendation of Gemeinholzer *et al*. (2010), Davis (2011), Culley (2013), Gostel *et al*. (2016) and Funk et al. (2017) regarding recording the presence of water streams, the soil type or the habitat (Table 3). Regarding sampling strategies, we found three categories in which our samples from the University of Oviedo collection can be classified: (1) taxonomic sampling, consisting in one or few individuals from various taxa sometimes including type locations (e.g. *Rivasmartinezia vazquezii* Fern.Prieto & Cires (2014), *Micranthes stellaris* (L.) Galasso, Banfi & Soldano (2005)), (2) genetic diversity sampling, comprising several individuals from the same taxa belonging to various populations (e.g. *Cochlearia pyrenaica* DC. (1821), *Ranunculus cabrerensis* Rothm. (1934)) and (3) distribution validation sampling, consisting in numerous individuals from one or more taxa throughout a wide geographical range (e.g. *Cytisus dieckii* (Lange) Fern.Prieto, Nava, Fern.Casado, M.Herrera, Bueno Sánchez, Sanna & Cires (2017), *Ulex cantabricus* Álv.Mart., Fern.Casado, Fern.Prieto, Nava & Vera (1988)). Hence, some taxa are represented by few individuals, while others count with dozens of individuals. Moreover, samples collected following a genetic diversity or a distribution validation sampling schemes are more likely to have associated ecological information than those obtained during taxonomic sampling. Besides, genetic diversity studies raw results have not been made publicly available in most cases, only the genetic parameters or statistical genetic analyses have been published, suggesting that a method of publishing these raw data should be developed to generate ESs and ESNs.

**Table 2.**
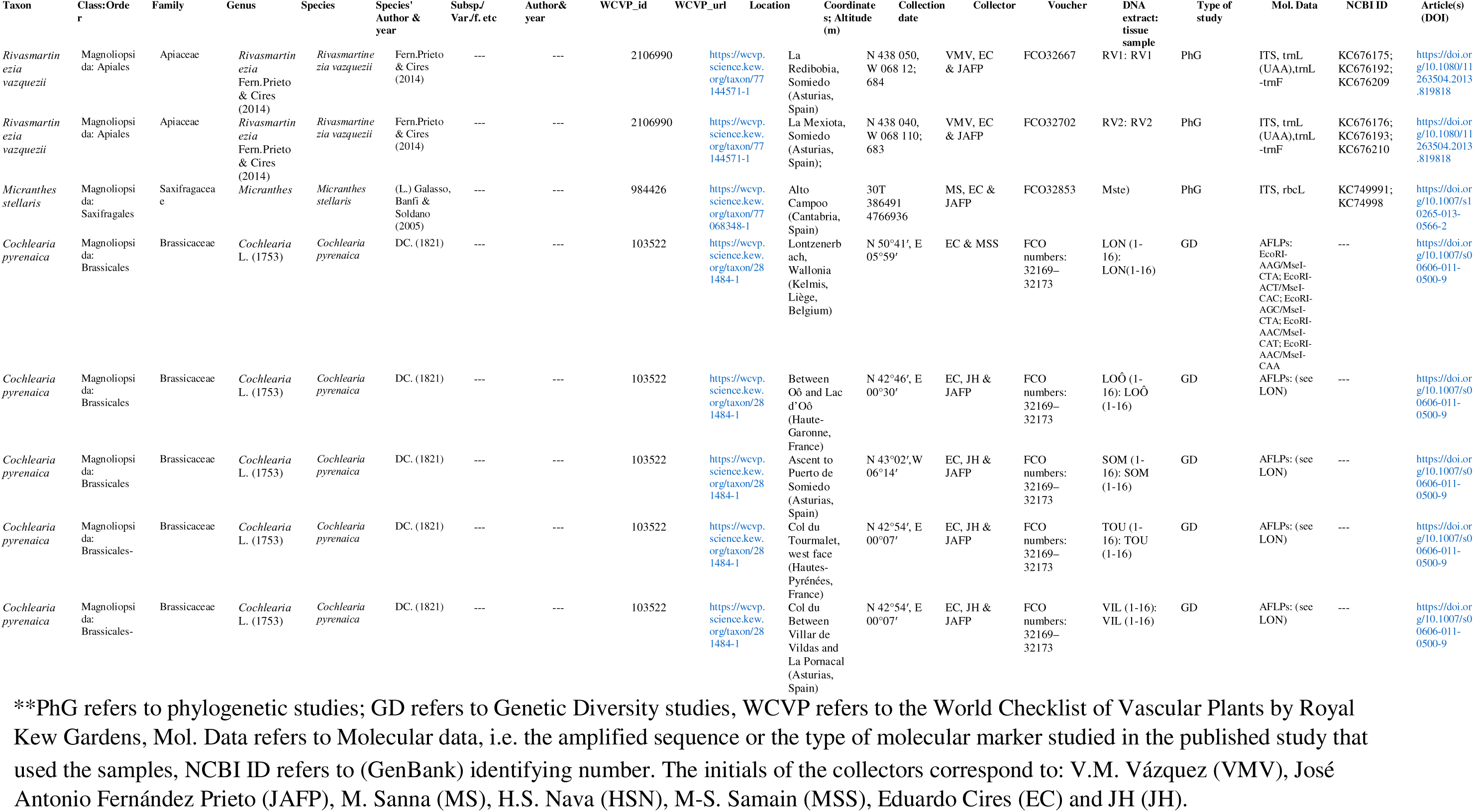
Summarised example of the data gathered from each of the DNA extracts and/or tissue samples of the 1182 collections of the area of Botany of the department of Biology of Organisms and Systems of the University of Oviedo.

**Table 3.**
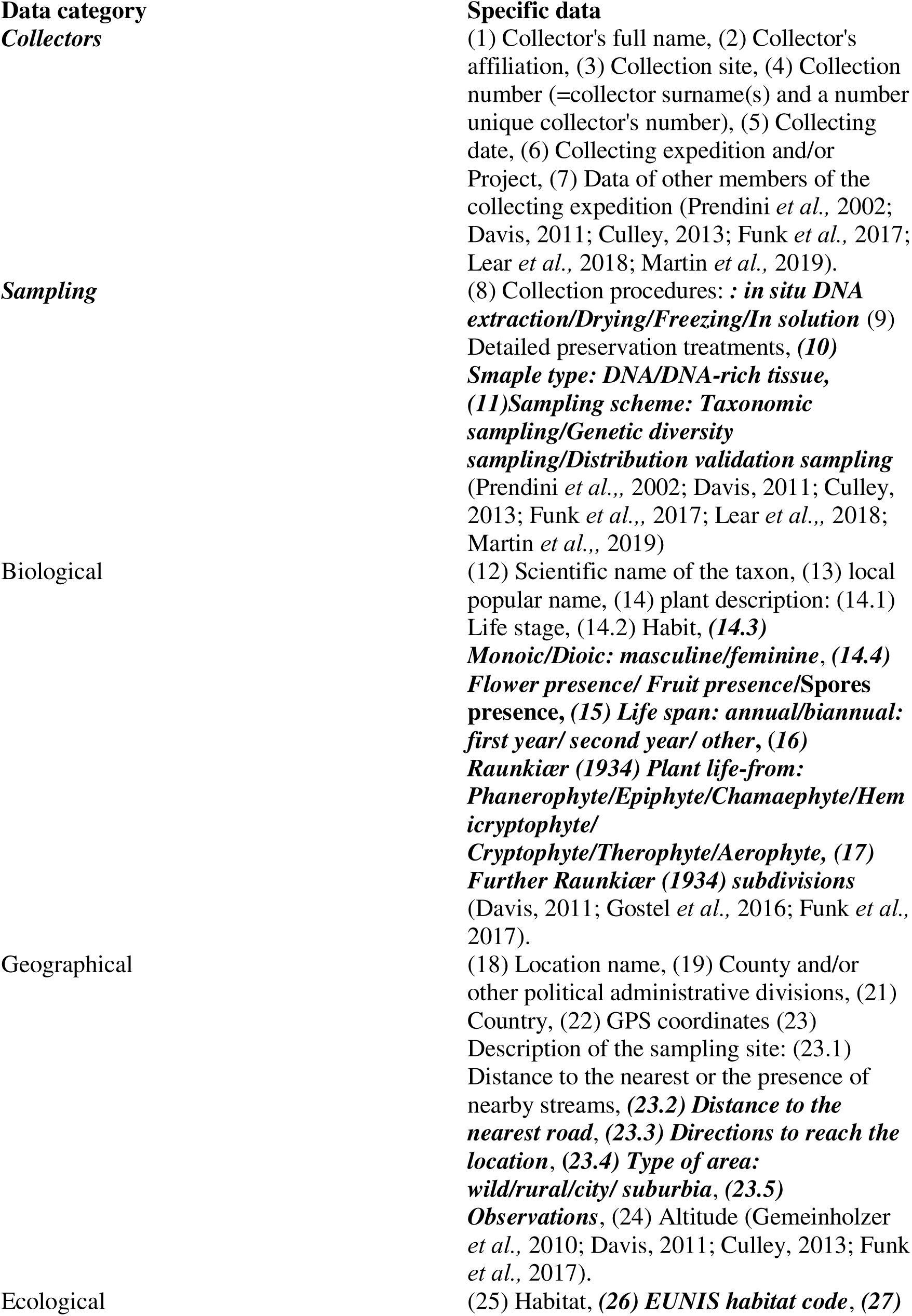
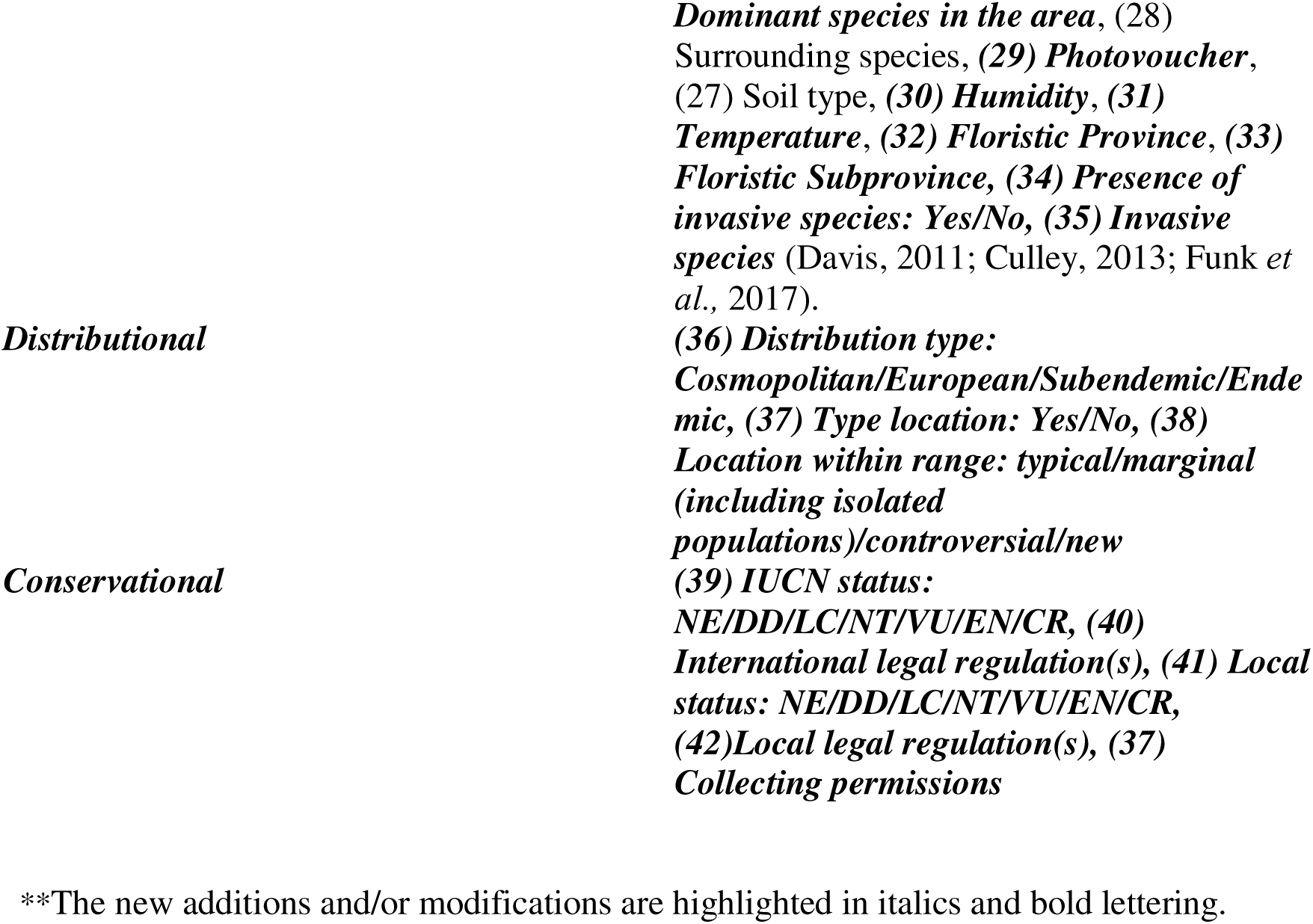
Modifications of the main categories of collection data proposed by previous authors and additions to the specific information.

### Phase 2: Determining the specific aims and the target area of the PBB

Given the current conservational necessities and the biodiversity loss projections for the next decades (De Vos *et al.,* 2015; Nic Lughadha *et al.,* 2020), establishing target areas and conservational aims should be a keystone for PBBs. Consequently, besides inventorying the already collected taxa, efforts should focus on comprehensively cataloguing the target area flora to determine conservational and scientific priorities. We classified these taxonomic, phylogenetic and conservational priorities in six groups: (1) target area endemisms, (2) subendemisms shared with other floristic regions, (3) putative endemic taxa with non-molecularly validated taxonomic status, (4) relictic and/or isolated populations, (5) taxa at conservational global risk and (6) taxa at conservational local risk. Furthermore, a detailed inventory could unveil understudied and/or undersampled taxa and areas, which would be relevant for future conservation strategies in collaboration with other biobanks.

Invasive Alien Species (IAS) are capable of causing a negative impact on biodiversity, socioeconomic activities and human health during their establishment and dissemination outside their native range (Kolar & Lodge, 2001; Keller *et al.,* 2011; Ricciardi, 2013).

The IASs presence throughout the planet is intimately related to human activities such as international commerce and to some features like their capacity of modifying nutrient cycles and soil features, which explains the fact that they pose the second most important threat to biodiversity (Westphal *et al.,* 2008; CBD, 2000; Vilà *et al.,* 2011; Ricciardi, 2013; Castro-Díez *et al.,* 2019; Shabani *et al.,* 2020). IASs genetic material collection at various locations within invaded areas will be of aid in understanding biological invasions features: (1) number of introduction events, (2) colonization waves potential origins, (3) temporal and spatial invasion dynamics, (4) predominant reproduction type (i. e. vegetative vs. sexual) and (5) genetic diversity difference between their invasive and natural ranges. Consequently, IASs sampling should be part of the PBB conservational priorities, given the genetic diversity’s crucial role in understanding and monitoring IAS spread.

### Phase 3: PBB structuring and setting the SOPs of the MOAs

DNA extracts and DNA-rich tissue collections can be organised into the recommended two-node PBB using the created inventories: DNA extracts collection would constitute the WN and the tissue collection would form the RN (Adams & Adams, 1992; Mattick *et al.,* 1992; Adams, 1997; Rice, Henry *et al.,* 2006; Hodkinson *et al.,* 2007; Zimkus & Ford, 2014a, 2014b; ISBER, 2018). Thus, DNA and tissue banking would be the first MOAs *sensu* Hodkinson *et al*. (2007) to have their SOPs set. Reasonable high-quality DNA preservation at low costs could be achieved by preserving DNA extracts at -20°C and DNA-rich tissues at room temperature in silica gel (Prendini *et al.,* 2002; Corthals & Desalle, 2005; Hodkinson *et al.,* 2007; Zimkus & Ford, 2014a, 2014b). Building the ESs collection would require high levels of traceability that could be achieved by implementing the Funk *et al*. (2017) double identifying number scheme, in which each sample has two identifying numbers: a primary identifying number derived from the voucher collecting number that connects ES samples and a second DNA Bank collection identifying number unique to each sample (Figure 4) (Caujapé-Castells *et al.,* 2006; ISBER, 2018). At this early stage, the labelling system should also follow the loaning samples recommendations of international organization (e. g. World Health Organization (WHO), 2003; ISBER, 2018; Instituto de Investigación de Recursos Biológicos Alexander von Humboldt (IPT IAvH), 2020).

**Figure 4.**
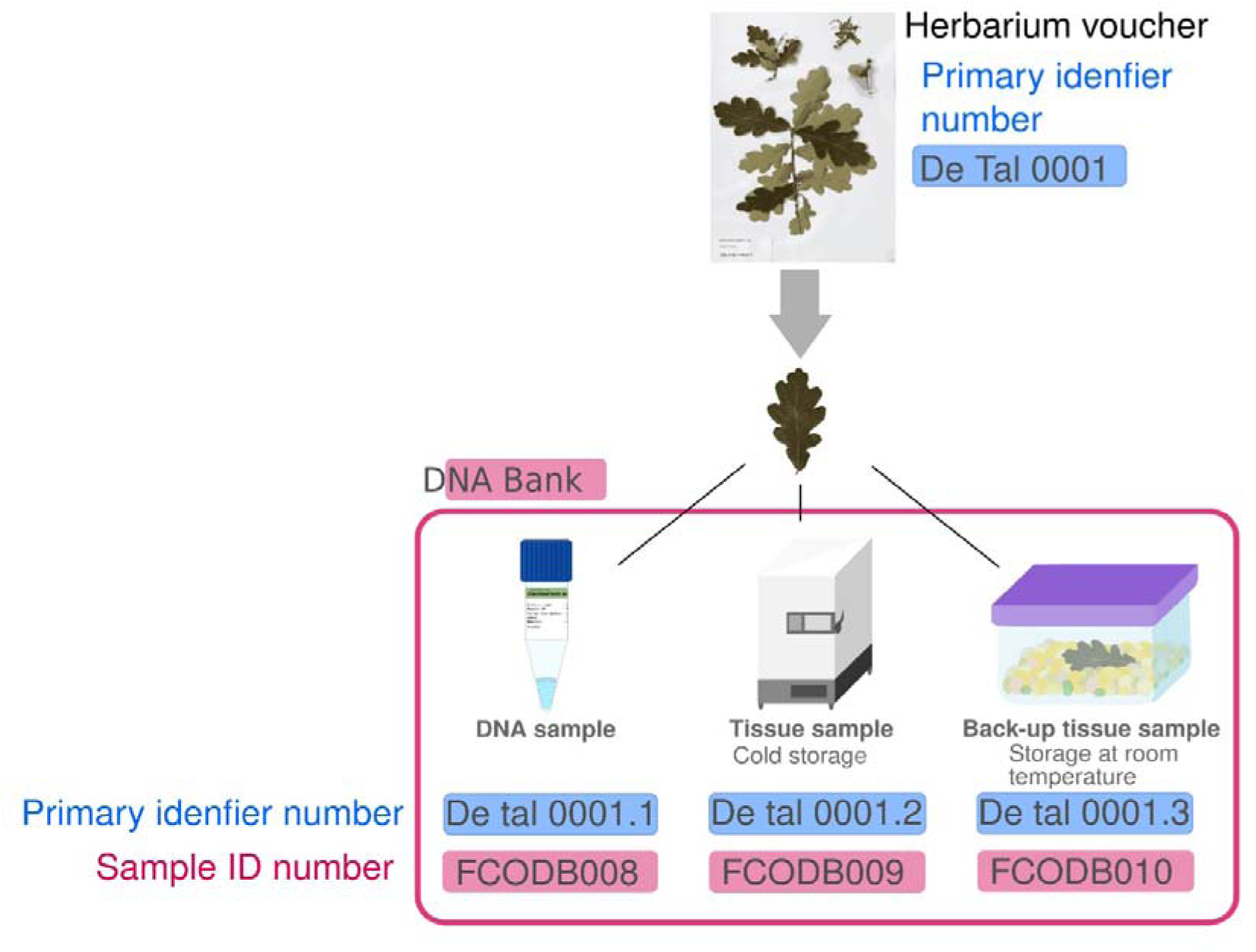
Scheme depicting an example of the Funk *et al.,* (2017) scheme for the system of double identifying number. The collecting number of a voucher used as primary identifier number of all the derived vouchers and the sample ID numbers of all tissues in DNA bank storage derived from the same sample. The collection number or primary identifier number connects the voucher and the samples since it is formed by the collecting number (in this case De Tal 00001) plus an additional number (e.g. De Tal 00001.1). When acquired by the DNA bank, another unique internal number, the sample ID number, analog to the Herbarium catalog number, is given to each of sample (in this case FCOD008).

A preliminary label design for the ESs collection should include: (1) product type (DNA, RNA or tissue), (2) collection number (=primary identifying number), (3) biobank ID number, (4) accepted genus, (5) accepted species, (6) sample origin and (7) an assigned barcode or QR code (WHO, 2003; Caujapé-Castells *et al.,* 2006; Campbell *et al.,* 2018; IPT IAvH, 2020) (Figure 5). Following the double identifying number system (Caujapé-Castells *et al.,* 2006; Funk *et al.,* 2017) and taking into account that at this stage there exist samples obtained before and after collection centralization, three different labels could be used: (1) the labels of samples collected before the centralization (i. e. initial ESs), (2) the collection labels (i. e. samples adquired after the SOPs were set) and (3) the loan labels. These three label types should be easily recognised (e.g. by using different colours or patterns) to guarantee sample’s traceability.

**Figure 5.**
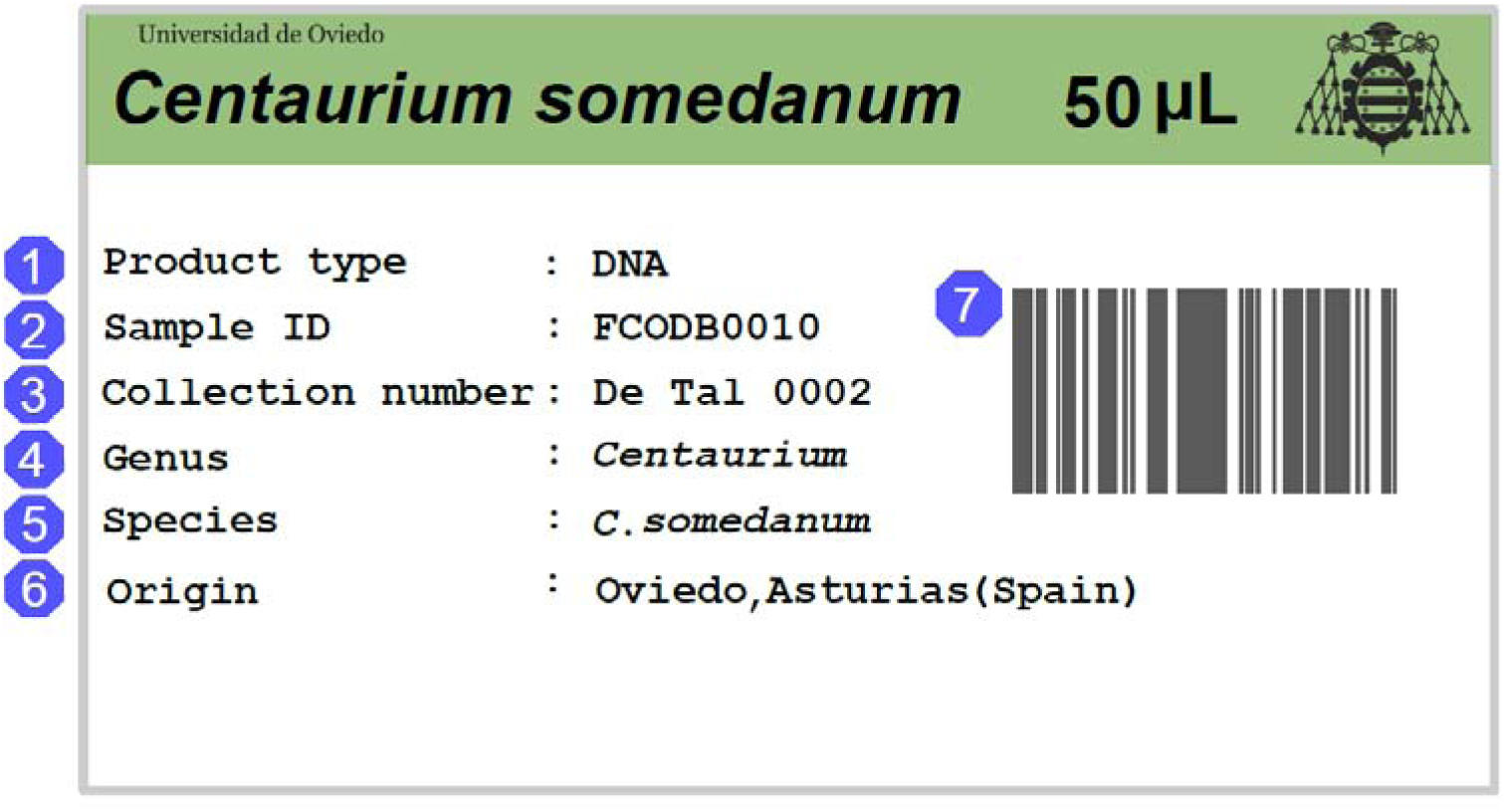
Example of potential label of loaned samples following the instructions of Wolrd Health Organization (WHO) (2003), Caujapé-Castells *et al.,* (2010), Campbell *et al.,* (2018a) and Instituto de Investigación de Recursos Biológicos Alexander von Humboldt (IPT IAvH), (2020). This authors believe that the label of a loaned sample should specify: the type of product (DNA, RNA or tissue) (1), the sample ID number of the biobank (2), the collection number (3), the genus of the sample (4), the species of the sample (5), the origin of the sample (6) and a barcode (7

Establishing the aims and priorities and the three collections interconnection (phases 1 and 2) would benefit material collection, data gathering and sample labelling SOPs by highlighting new ways of improving ESs and ESNs to match the scientific necessities. For example, updating the Flora checklist brings the opportunity to centralize ecological (e.g. European Nature Information System (EUNIS) habitats (https://eunis.eea.europa.eu/), IUCN Habitats Classification Scheme (https://www.iucnredlist.org/resources/habitat-classification-scheme), or floristic subprovince), distributional (e.g. endemisms, subendemisms or isolated populations) and conservational data (e.g. IUCN (https://iucn.org/) assessments or local assessments); while implementing local legislation and the Protocol of Nagoya. As we believe this information should be part of the SOPs related to ESs and ESNs, we propose to combine and extend the data collection items suggested by previous authors (e. g. Prendini *et al.,* 2002; Culley, 2013; Lear *et al.,* 2018) to specify: (1) national and international conservational status, (2) sampling location features regarding distribution range, (3) the sampling scheme and (4) habitat features and classification (Table 2).

At a second stage, sample collection and DNA extraction and utilization SOPs could be established, implementing legal documents for donations, collections and loans while abiding regional, national and international legislations (e. g. the Protocol of Nagoya) (Davis & Borisenko, 2017; CETAF, 2020). Thus, enforcing the ABS agreements by establishing the guidelines to generate PICs, MATs and MTAs, which are essential to sample collection and DNA utilization (CBD, 2010; GGBN, 2015; Biber-Klemm *et al.,* 2016; Davis & Borisenko, 2017). International organizations such as the Middle Eastern and African Society for Biopreservation and Biobanking (ESSB) (https://esbb.org/), the Consortium of European Taxonomic Facilities (CETAF) (https://cetaf.org/) or the International Society for Biological and Environmental Repositories (ISBER) (https://www.isber.org/) can facilitate this thanks to their remarkable contributions to DNA biobanking like producing documents for the ABS agreements implementation and especially DNA samples MTAs (e.g. GGBN, 2015; CETAF, 2020) or the ISBER Best Practices Guidelines (ISBER, 2018).

### Phase 4: Go public: from three collections to ESs and ESNs

Once the SOPs are established and the three collections connected, a website should be built with the following characteristics: (1) be user-friendly, (2) interconnect specimens’ data generating ESs and ESNs and (3) be informative regarding the biobank functioning, samples’ availability and the donation/request process. The best practice guides and ABS agreements guidelines generated during phase 4, would be of capital importance for the informative and user-friendly aspects. Furthermore, generating ESs and ESNs from herbarium voucher would upgrade the ongoing herbarium digitalization processes by producing and fomenting FAIR data and the RRI (Owen *et al.,* 2012; Wilkinson *et al.,* 2016; Lannom *et al.,* 2019). Opening the website to the public would be the final step of the three collections interconnection.

### Phase 5: Becoming part of GGBN and other organizations and establishing collaborations

Once the PBB is functioning, becoming part of the GGBN should be the next step as this is opened to recruiting new members and aims to standarise the MOAs (Dröge *et al.,* 2014; Seberg *et al.,* 2016; GGBN, 2024). This membership would improve the PBB visilibity among reasearchers and encourage collaborations other biorepositories.

During this phase, the PBB could also aim to become part of other associations like ESSB (https://esbb.org/), CETAF (https://cetaf.org/) or ISBER (https://www.isber.org/).

Relocating the RN in other countries or harbouring samples from other biobanks was suggested during the early days of biodiversity DNA biobanking (Adams, 1997; Graner *et al.,* 2006; SANBI, 2020) and has been implemented by some PBBs like the SANBI DNA bank (https://www.sanbi.org/biodiversity/foundations/genetics-services/) and the Kew DNA Bank (https://dnabank.science.kew.org/homepage.html)). While other PBBs are willing to do so (e. g. Jardín Botánico Canario “Viera y Clavijo” (http://www.jardincanario.org/banco-de-adn-motivacion)). In this sense, sharing and exchanging samples would benefit both biobanks, especially when both share floristic elements. For instance, two different PBBs could harbour samples of closely related taxa occurring in different bioclimatic regions or of relictic and non-relictic populations or samples from non-native taxa at locations where they behave as IASs and at locations where they are innocuous species. In this way, active interactions and partnerships with other biobanks have the potential of boosting research in fields like biogeography, conservation biology or taxonomy.

### Phase 6: Keep improving

Since high-quality DNA preservation is a key aspect of biobanking, the PBB should focus some of its research activity on finding and assessing new preservation methods to lower costs and risks. The performances of different solutions allowing storing DNA at room temperature (Wan *et al.,* 2010; Clermont *et al.,* 2014) or DNA extraction methods applied to different plant groups could be assessed, making methodological contributions to the scientific community. Similarly, new ways of improving ESs and ESNs should an aim of PBBs.

## Discussion

The BDC—one of the main ecosystemic, societal and economic threats to humanity— is characterised by extinction rates 1000 times higher than expected and has been exacerbated during the past decades by factors such as Climate Change, habitat loss and fragmentation or IAS; leading to predictions of reaching 10000 times the background extinction rate (Feeley & Silman, 2010; Dullinger *et al.,* 2012; De Vos *et al.,* 2015; Ceballos *et al.,* 2017; Muthukrishnan & Larkin, 2020; Shabani *et al.,* 2020; Bellard *et al.,* 2021; Eichenberg *et al.,* 2021; Prakash & Verma, 2022; Pörtner *et al.,* 2023; Williams *et al.,* 2023). The scientific community addressed this threat by generating and making available new phylogenetic knowledge, while national administrations and international organizations set different aims and funded biodiversity conservation programs like the Global Strategy for Plant Conservation (GSPC) of the United Nations (Blackmore, 2002; United Nations (UN), 2021). All these efforts have not decelerated plant species decline and extinctions, which can have unpredictable and disproportioned effect on ecosystems (Grime, 2002; Schleuning *et al.,* 2016; Eichenberg *et al.,* 2021).

More than 170 plant species have gone extinct in the wild since the 1960s, around 50,000-75,000 may currently be at risk of extinction and many could go extinct before even been described (Cronk 2016; Pimm & Raven, 2017; Willis, 2017; Ruta *et al.,* 2020; IUCN, 2024). Moreover, the prevention of local and global plant extinctions faces various additional problems: (1) chronical lack of funding (Suarez & Tsutsui, 2004; Balding & Williams, 2016; Roberson & Meyer, 2020), (2) lack of time for sampling the declining species and populations due to the synergetic interactions between extinction- driving factors (e .g. climate change and IAS), (4) absence of unified species conservation and IAS erradication strategies sometimes even within the same country (e. g. *Acacia mearnsii* De Wild. (1925), *Cortaderia selloana* (Schult. & Schult.f.) Asch. & Graebn. (1900)), (5) inefficient or incomplete systems of information sharing (i. e. there is not a conservation strategies database that facilitates decision-making based on previous experiences), and (6) the reduced percentage of species with genetic information (Nic Lughadha *et al.,* 2020).

Species and populations genetic data is as relevant for plant species conservation as habitat restauration and landscape dispersal securing (Aavik & Helm, 2018). Centralizing and systematizing Department DNA collections would be of aid in implementing the genetic information use in conservation programs and would also improve IUCN assessments’ accuracy (Garner *et al.,* 2020). Furthermore, the Dynamo scheme, by connecting smaller collections, could boost collaborations in genetic diversity studies and conservation efforts as different collections could contribute by sampling local populations, thus covering a larger proportion of the distribution range. Moreover, prioritarian taxa inventories would encourage conservational information sharing, including IAS dynamics and erradication strategies. In this context, more efficient conservation plans and the ABS agreements implementation (CBD, 2010) would also benefit society by preserving and regulating the plant genetic and biodiversity heritage, while addressing the GSPC goals (UN, 2021).

During the last decade, iniciatives of international organizations like the GGBN’s Global Genome Initiative-Gardens (GGI-Gardens) (https://naturalhistory.si.edu/research/global-genome-initiative), the Biodiversity Collections Network (BCoN) (https://bcon.aibs.org/) or the Distributed System of Scientific Collections (DiSSCO) (https://www.dissco.eu/what-is-dissco/the-collections) have contributed to natural collections digitalization and to completion of the vascular plants genetic barcoding (Gostel *et al.,* 2016; Zúñiga *et al.,* 2017). In the actual context, generating ESs collections and ESNs databases and providing updated inventories at a global scale could be viewed as the next logical step to these digitalization initiatives. Nevertheless, the proposed initial steps to this process, like creating a common database similar to the Index Herbariorum (http://sweetgum.nybg.org/science/ih/) listing the existing PBBs, have not been implemented at a global scale (Spooner & Ruess, 2014a), although the USA chieved this through the Integrated Digitized Biocollections (https://www.idigbio.org/).

Our 6-step plan for transforming 3 separate collections into a ESs and ESNs PBB would benefit several scientific fields—e. g. molecular taxonomy, conservational biology, DNA barcoding or eDNA—thanks to its positive impact on research results reproducibility and reliability (Corthals & Desalle, 2005; Rice, Henry *et al.,* 2006; Astrin *et al.,* 2013; Spooner & Ruess, 2014b; De Vere *et al.,* 2016; Lear *et al.,* 2018).

This process represents additional advantages for local and regional conservational and research goals. For instance, our collection focuses on the Cantabrian Mountains flora (northern Iberian Peninsula), establishing an small PBB would imply the application FAIR data and RRI principals and achieving some the goals for research improvement and sustainability of European Union’s (EU) Horizon Europe (Owen *et al.,* 2012; Wilkinson *et al.,* 2016; Lannom *et al.,* 2019; European Union (EU), 2021), while giving a new dimension to the EU initiative Innovation and Consolidation for large scale Digitisation of natural heritage (ICEDIG) (European Commission, 2023) by generating ESNs (e. g. connecting herbarium voucher occurrences with their genetic data).

Moreover, PBBs would make the goals of international barcoding projects such as the International Barcode of Life Project (iBOL) (https://ibol.org/), Encyclopedia of Life (EOL) (https://eol.org/) or The Barcode of Life Data System (BOLD) (https://www.boldsystems.org/) easier to achieve, while other fields like eDNA, SNPs genotyping, transcriptomics or phylogenomics would enhance their accuracy (Adams, 1997; Rice, Shepherd *et al.,* 2006; Hanner & Gregory, 2007; Pleijel *et al.,* 2008; Rice, Kasem & Henry, 2008; Astrin *et al.,* 2013; Spooner & Ruess, 2014b; De Vere *et al.,* 2016).

Our 6-step process was designed to improve our knowledge on the relatively unstudied local Cantabrian Mountains vascular flora through the generation of a database and an updated inventory (Izco, 1980; Fernández Prieto & Vázquez, 2007; Fernández Prieto *et al.,* 2014; Gómez Durán, 2014), aiming to unveil taxonomic knowledge gaps and the factors associated to local biodiversity hotspots. With this information, the promotion of large expeditions focused on understudied areas and taxonomic groups would simultaneously contribute to the Leipzig Catalogue of Vascular Plants and tackle one of the major bottlenecks for genetics-based studies (Mattick *et al*., 1992; Lear *et al.,* 2018; SANBI, 2020; Freiberg *et al.,* 2020). Particularly, ecology and molecular taxonomy would potentially profit from an updated species catalogue thanks to data accuracy refinement and the possibility of sampling the type localities of taxa with scientific interest. In this sense, University of Oviedo department collections holds a very good example of the relevance of implementing DNA sampling when describing new taxa.

Our collections includes specimens from the only two known species of the subendemic genus *Rivasmartinezia* Fern.Prieto & Cires (2014)— *Rivasmartinezia vazquezii* Fern.Prieto & Cires (2014) and *Rivasmartinezia cazorlana* Blanca, Cueto, Benavente & J.Fuentes (2016)— collected from their type localities (Fernández Prieto & Cires, 2013; Blanca *et al.,* 2016). Consequently, we could provide new molecular information (e. g. sequencing new different molecular markers) to the already available data at GenBank extending de facto ESs and ESNs without resampling. In this way, many small department PBBs connected through the Dynamo scheme could enhance information sharing by generating and expanding ESs and ESNs, even when they are type specimens. Furthermore, international digitalization and biodiversity proyects would progress more rapidly in our biodiversity loss contexts if there existed many small department PBBs able to provide this type of specimens following standardized protocols.

## Acknowledgements

Claudia González-Toral had the initial financial support of the Government of Asturias (2002166- Programa Severo Ochoa) and is currently supported by the Ramón Areces Foundation PostDoctoral Fellowship program (XXXVI Convocatoria para Ampliación de Estudios en el Extranjero en Ciencias de la Vida y de la Materia). The authors would also like to thank Estrella Alfaro-Saiz, Juan Arroyo, Juan Viruel, Pablo Muñoz, Herminio S. Nava and Candela Cuesta for their useful comments and advice.

## Data Accessibility Statement

All data used for this review has been exposed in the text.

## Funding statement

Claudia González-Toral had the initial financial support of the Government of Asturias (2002166- Programa Severo Ochoa) and is currently supported by the Ramón Areces Foundation PostDoctoral Fellowship program (XXXVI Convocatoria para Ampliación de Estudios en el Extranjero en Ciencias de la Vida y de la Materia).

## Conflict of interest disclosure

The authors have no conflicts of interest to disclose.

## Notes

### Competing Interest Statement

The authors have declared no competing interest.

